# Comparing fMRI responses measured at 3 versus 7 Tesla across human cortex, striatum, and brainstem

**DOI:** 10.1101/2020.05.12.090175

**Authors:** Olympia Colizoli, Jan Willem de Gee, Wietske van der Zwaag, Tobias H. Donner

## Abstract

Significant progress has been made in ultra-high field functional magnetic resonance imaging (fMRI) at 7 Tesla (T). While fMRI at 7 T promises a general increase in sensitivity compared to lower field strengths, the benefits may be most pronounced for specific applications. The current study aimed to evaluate the relative benefit of 7 T over 3 T fMRI for the assessment of task-evoked fMRI responses in different brain regions. We compared the amplitude of task-evoked responses between 3 T and 7 T measured from the same human participants. Participants performed a challenging random dot motion discrimination task with delayed monetary feedback, which animal physiology has linked to several cortical and subcortical structures including extrastriate (dorsal) visual cortical areas, the striatum, and the brainstem including dopaminergic midbrain nuclei. We quantified the evoked fMRI responses in each of these brain regions during the decision interval and the post-feedback interval of the task, and compared them between brain regions and field strengths. The dependence of response amplitudes on field strength during the decision interval differed between cortical, striatal, and brainstem regions, with a generally bigger 7 T vs. 3 T benefit in subcortical (in particular brainstem) structures. We also found stronger differential responses during easy than hard decisions at 7 T for the dopaminergic nuclei, possibly reflecting reward expectation. Our results demonstrate the potential of 7 T fMRI for illuminating the contribution of small brainstem nuclei to the orchestration of cognitive computations in the human brain.

**Highlights:**

- We compared 7 T to 3 T fMRI during perceptual decision-making under uncertainty.
- Differences between 7 T and 3 T evoked responses and tSNR varied across the brain.
- Evoked responses in dopaminergic brainstem nuclei were bigger at 7 T than 3 T.
- The responses of dopaminergic nuclei are consistent with reward expectation.
- Results highlight the potential of 7 T fMRI for imaging small brainstem nuclei.

## 1. Introduction

The last decade has shown significant progress in ultra-high field (UHF) Magnetic Resonance Imaging (MRI), providing excellent image quality and sub-millimeter resolution for functional, structural and connectivity images. 7 T MRI is on its way to becoming a user-friendly modality, with sequence development centered around increasing spatial and temporal resolution, as well as improving the spatial specificity of the blood oxygenation level dependent (BOLD) signal for studies of brain function (for a review see (van der Zwaag et al., 2016)). Accordingly, 7 T functional MRI (fMRI) has been progressively used for studying brain functions including perception, motor action, decision-making, language, and emotion in normal subjects (e.g. (Bode et al., 2011; Harvey et al., 2013; Hayashi et al., 2018; Mestres-Missé et al., 2012; Theysohn et al., 2013; Thürling et al., 2012, 2011; Torrisi et al., 2018; van der Zwaag et al., 2012). Given the increasing availability of 7 T MRI scanners, an important question facing researchers pertains to which fMRI applications in particular benefit most from 7 T fMRI, compared to more standard (cheaper and more widely available) measurements at 3 T.

In general, 7 T is expected to improve the image signal-to-noise ratio (SNR) of MRI measurements relative to standard 3 T acquisitions (Edelstein et al., 1986; Pohmann et al., 2016). For BOLD fMRI this predicts stronger evoked fMRI responses (Turner et al., 1993; van der Zwaag et al., 2009; Yacoub et al., 2001). Yet, increased field strength is also accompanied by an increase in susceptibility-induced distortions, such as at air-water interfaces and the contributions from physiological noise (Triantafyllou et al., 2005; van der Zwaag et al., 2016). Physiological (cardiac and respiratory) noise is greater in the brainstem than in the cortex (Harvey et al., 2008). Given the above-described trade-offs, fMRI at 7 T is not necessarily always preferable to fMRI at 3 T.

Previous studies have used simple blocked designs with visual or motor tasks to assess the advantages of 7 T for BOLD-fMRI measurements (e.g. (Duong et al., 2002; Schäfer et al., 2007; van der Zwaag et al., 2009; Yacoub et al., 2003, 2001). Other studies have compared 7 T and 3 T fMRI for clinical applications (e.g. (Beisteiner et al., 2011; Morris et al., 2019) or investigated task-evoked activation related to cognition (de Hollander et al., 2017; Theysohn et al., 2013; Torrisi et al., 2018). Yet, only some of these studies have been direct head-to-head comparisons between functional measurements at the two field strengths in the same individuals. Furthermore, these studies have either focused on the cerebral cortex or did not explicitly discriminate between cortical and subcortical brain regions in the field strength comparisons.

Here, we aimed to fill those gaps, focusing on the brainstem. The brainstem is a part of the brain for which fMRI studies may substantially benefit from 7 T. The brainstem is made up of a multitude of small nuclei, precise delineation of which requires higher spatial resolution than for cortical or even other subcortical regions (e.g. the striatum). Despite successful attempts using sophisticated measurement protocols (D’Ardenne et al., 2008; de Gee et al., 2017; Iglesias et al., 2013), the brainstem is notoriously difficult to image with 3 T fMRI. At the same time, the brainstem is an important target for cognitive (de Gee et al., 2017; Iglesias et al., 2013) and translational (Stephan et al., 2015) neuroimaging. This is because the brainstem houses the nuclei of the brain’s so-called neuromodulatory (e.g., dopamine or noradrenaline) systems, which send widespread, ascending projections to higher parts of the brain including the cerebral cortex, and seem to play a key role in cognition and disturbances thereof (Aston-Jones and Cohen, 2005; Dayan, 2012; Montague et al., 2004).

The current study evaluated the relative benefit of 7 T over 3 T fMRI for the assessment of task-evoked fMRI responses across a range of select brain regions, including motion-responsive visual cortical regions and anterior cingulate cortex, the striatum, and the noradrenergic locus coeruleus (LC) as well as dopaminergic ventral tegmental area (VTA) and substantia nigra (SN). To this end, we used a direct head-to-head analysis of task-evoked fMRI responses as well as the temporal SNR (tSNR) of ongoing signal fluctuations within the same healthy adult participants scanned at 3 T and 7 T. The task was designed so as to enable the assessment of different components of task-evoked neural responses (in different trial intervals), the neurophysiological underpinnings of which have been well-characterized in the above structures by previous single-unit neurophysiology in rodents and monkeys.

## 2. Methods

We acquired fMRI (near-whole brain coverage) and behavioral data in a 3 T and a 7 T setup with the same healthy adult volunteers who performed a two-alternative forced choice (2AFC) motion-discrimination task with monetary-coupled feedback. Independent analyses of the relationship of choice behavior and participants’ pupil responses during the 3 T sessions have been published previously (Colizoli et al., 2018; de Gee et al., 2017). The analyses presented in the current paper are conceptually and methodologically distinct. Relevant methods that have previously been published are again summarized in section 2.

### 2.1. Participants

All participants were screened for MRI-related health risks using a standard procedure and gave written informed consent. The experiment was approved by the Ethical Committee of the Department of Psychology at the University of Amsterdam. Participants were financially compensated with 10 Euros per hour in the behavioral lab and 15 Euros per hour for MRI scanning. In addition to this standard compensation, participants earned money on the 2AFC task based on their performance within each scan session: €0–10 linearly spaced from 50– 100% accuracy for each scan session (i.e. 50% correct = €0, 75% = €5, 100% = €10). At the end of each 2AFC run (25 trials, see section 2.3), participants were shown (on screen) their average performance accuracy across the runs in the current session and their corresponding monetary reward.

Ten participants (two authors) completed both the 3 T and 7 T MRI measurements. Due to technical failure, physiological recordings were lost during the 7 T acquisition for more than 50% of the data of two participants and for six individual runs (across three participants). These two participants were, therefore, excluded from further analysis. This yielded a sample of eight participants (including one author; 3 women, *M* = 28 y, *SD* = 4.8, range 23-37), in which we performed head-to-head comparison reported in this paper.

The 3 T acquisition data from one session of one participant were excluded from further analysis because of motion-related MRI artifacts.

### 2.2. Procedure

Each participant completed one behavioral session to determine individual 70% and 85% correct motion-discrimination thresholds (in terms of motion coherence; see section 2.3) before completing several MRI sessions across different days. During the 3 T and 7 T MRI acquisitions, participants lay supine in the scanner, and their heads were secured with additional padding to minimize motion. Participants wore ear protection and headphones and were instructed to maintain a central fixation during all functional runs. The cardiac cycle was monitored with a pulse oximeter attached to the left ring finger. Respiratory activity was recorded with a chest belt. Physiological signals were recorded at a sampling rate of 496 Hz. Stimuli were presented on a 31.55” MRI compatible LCD display with a spatial resolution of 1920×1080 pixels and a refresh rate of 120 Hz.

#### 3 T MRI experiment

Participants viewed the projection screen from 120 cm via a mirror attached to the head coil. Sessions began with reference and anatomical scans, during which participants practiced the 2AFC motion-discrimination task (1-2 blocks of 50 trials). Participants were administered six runs of the 2AFC task during each scan session. After the sixth run, a B0-field map was acquired. A coherent-motion localizer was administered at the end of the scan session (section 2.10). Each session took 2 hours. Participants completed four sessions across different days.

#### 7 T MRI experiment

Participants viewed the projection screen from 208 cm via a mirror attached to the head coil. Sessions began with reference scans, during which participants practiced the 2AFC motion-discrimination task (1-2 blocks of 50 trials). Participants were administered six runs of the 2AFC task during each scan session. After the sixth run, a B0-field map was acquired. A coherent-motion localizer was administered at the end of the scan session. Each session took 1.5 hours. Participants completed two sessions across different days.

### 2.3. 2AFC motion-discrimination task

Participants were asked to decide whether the net direction of motion of a dynamic random dot kinematogram (presented for 750 ms) was upwards or downwards; the ratio of coherently moving ‘signal dots’ to incoherently moving ‘noise dots’ determines the difficulty level on each trial (hard or easy, 70% vs. 85% correct, respectively). The motion coherence levels corresponding to these two difficulty (i.e., performance) levels were individually determined using a staircase procedure with seven levels (100 trials per level, 50 trials per visual hemifield, levels were: 0%, 2.5%, 5%, 10%, 20%, 40% and 80% coherence) in a separate behavioral session that took place before the MRI sessions. During the 2AFC task, motion coherence varied randomly from trial-to-trial so that observers performed at 70% correct in 2/3 of trials (‘hard’) and at 85% correct in 1/3 of trials (‘easy’). After a variable delay following the choice on each trial (3.5–11.5 s, uniformly distributed across 5 levels, steps of 2 s), feedback was presented (green fixation square for correct, red for error, 0.42 s) that was coupled to a monetary reward (see section 2.2 for details). The inter-trial interval lasted 3.5–11.5 s, uniformly distributed across 5 levels, steps of 2 s. During all phases of a trial, with the exception of the stimulus presentation of coherent motion, random dot motion (0% coherence) was presented.

All stimuli were presented within a central annulus (not visible to participants). The annulus contained 524 dots all within one visual hemifield (left or right; counterbalanced across runs and participants). The different screen viewing distances between the two scanner environments (see above) caused some differences in the visual stimulation parameters: The annulus outer diameter was 16.8° or 9.4° and dots moved at 7.5°/s or 3.9°/s during the 3 T or 7 T acquisitions, respectively.

One run of the 2AFC task contained 25 trials. Because subjects performed twice as many sessions at 3 T than at 7 T (see above) we split the 3 T data into two subsets (3 T-A and 3 T-B; see section 2.6) with 9-12 runs distributed over two sessions included in the analysis for each of the data subsets and, yielding a total of 225-300 trials per 3 T subset per participant. For the 7 T acquisition, 9-12 runs distributed over two sessions were included in the analysis, yielding a total of 225-300 trials per participant.

### 2.4. MRI data acquisition at 3 T

MRI data were acquired on a 3 T Philips Achieva XT MRI scanner (Amsterdam, the Netherlands) using a 32-channel dStream head coil across four sessions. Single-shot, 2D gradient echo-planar images (EPI) were acquired in 33 slices oriented perpendicular to the floor of the fourth ventricle and to the longitudinal extent of the locus coeruleus (Keren et al., 2009): 2 × 2 mm in-plane resolution, 3 mm slice thickness (no gaps), FOV 192 × 192 × 99 mm. TR = 2 s, TE = 27.62 ms, flip angle = 76.1°, SENSE acceleration factor = 3.0. A structural T1-weighted scan was acquired for anatomical co-registration and cortical surface reconstruction (voxel size: 1 × 1 × 1 mm^3^, TR = 8.13 ms, TE = 3.7 ms, flip angle = 8°). A B0 field map was acquired for unwarping (voxel size: 2 × 2 × 2 mm^3^, TR = 10.85 ms, TE = 3.03, flip angle = 8°). Slices were acquired sequentially in the ascending (foot to head) direction.

We chose anisotropic voxels and the above orientation of the imaging volume for EPI to optimize spatial sampling for the locus coeruleus (for details, see (de Gee et al., 2017)). The locus coeruleus is an elongated structure with small (a few millimeters) diameter and comparably large longitudinal extent (∼15 mm) along the floor of the fourth ventricle (Betts et al., 2019). The rationale was that the locus coeruleus was the smallest brainstem nucleus to be assessed here.

### 2.5. MRI data acquisition at 7 T

MRI data were acquired on a 7 T Philips Achieva MRI scanner (Amsterdam, the Netherlands) using a 32-channel Nova head coil across two sessions. Gradient EPIs were acquired in 30 slices oriented perpendicular to the floor of the fourth ventricle: 1.5 × 1.5 mm in-plane resolution, 3 mm (no gaps) slice thickness, FOV 192 × 192 × 90 mm. The voxel size was reduced in the in-plane direction compared to the 3 T acquisition, again because of the geometry of the locus coeruleus described in section 2.4 above. This way, the higher SNR at 7 T was traded for resolution. TR = 2 s, TE = 23 ms, flip angle = 70°, SENSE acceleration factor = 2.5. The TE and flip angle of the EPI differed from those in the 3 T protocol to accommodate for the shorter T_2_^*^ and longer T_1_ at 7 T. Finally, a B0 field map was acquired (voxel size: 2 × 2 × 2 mm^3^, TR = 4.6 ms, TE = 1.88, flip angle = 10°). Slices were acquired in the ascending (foot to head) direction in an interleaved fashion (first odd then even slices).

### 2.6. fMRI preprocessing

Pre-processing steps were identical for the 3 T and 7 T acquisitions. Each run of the EPI images was (i) brain extracted using the BET tool as part of FSL version 5.0.2.1 (Jenkinson et al., 2012; Smith et al., 2004; Woolrich et al., 2009), (ii) unwarped based on the session-specific B0 field map (FUGUE, FSL), (iii) motion corrected (sinc interpolation, MCFLIRT, FSL), and (iv) high-pass filtered to correct for low frequency drifts (Gaussian-weighted least-squares straight line fitting, window size = 50 samples). For each session, one EPI volume, the motion-corrected mean volume of the third run (out of six) of the 2AFC task, was used as the reference image for all EPI-based registration including motion correction. The reference image of each session was transformed to the T1-weighted MNI-152 2-mm space, via the subject-specific T1-weighted anatomical image, using an affine transformation with 12 degrees of freedom, the normmi cost function, and sinc interpolation (FLIRT, FSL). Within each session, all EPI runs were subsequently concatenated in time.

Nuisance regression was applied to the concatenated EPI runs for each session separately, including physiological noise correction in which cardiac and respiratory phases were assigned separately for each slice in the concatenated EPI time series (an extended version of RETROICOR (PNM, FSL)) (Brooks et al., 2013; Glover et al., 2000). Nuisance regressors included 34 physiological signal components and the mean signal fluctuation within the 4^th^ ventricle. The fourth ventricle was manually defined by marking voxels covering the ventricle on the basis of the T1-weighted MNI-152 2-mm space (52 voxels) and transformed to the subject- and session-specific 3D space (FLIRT). For the tSNR analysis, task effects in the data were removed by including two additional nuisance regressors: the task-evoked responses locked to the stimulus and feedback onsets convolved with a double gamma hemodynamic response function (HRF; 32 s kernel, peak amplitude of 1 at 6 s, undershoot amplitude 1/6th of response size at 16 s). The amplitude of the stimulus-locked response was modulated by reaction times (RT) on a trial-by-trial basis. The residuals of the nuisance regression were transformed to the T1-weighted MNI-152 2-mm space and used in the subsequent analyses. Whole brain analyses were restricted spatially by the common field of view between the two field-strength acquisitions.

We took advantage of the additional 3 T sessions (four) as compared with the 7 T sessions (two), by splitting the 3 T data set into two subsets (‘3 T-A’ & ‘3 T-B’) in order to compare the 7 T data set to a single 3 T data set (see sections 2.7 and 2.8).

### 2.7. Quantifying task-evoked BOLD responses

The pre-processed time series were entered into a general linear model (GLM) using FEAT version 6.0 (FSL v. 5.0.2.1). One GLM per participant was fit to each of their data sets (3 T-A, 3 T-B & 7 T). Session data were concatenated within each data set (1-2 sessions per data set per participant). The number of trials entered into each GLM was equalized across data sets for each participant according to the minimum of either the 3 T-A, 3 T-B or 7 T acquisitions. Regressors of interest were the following: two regressors for the responses during the decision interval (time-locked to onset of dot motion stimulus), for easy and hard trials, separately. The easy and hard stimulus regressors were parametrically modulated by RT. Two regressors modelled the feedback responses, for correct and error trials, separately. Nuisance regressors included one regressor for missed trials (stimulus and feedback responses were combined), and 1-2 regressors (each) modeling a session’s mean value. All regressors were convolved with the double-gamma HRF and high-pass filtered (cut-off at 100 s). Grand-mean intensity normalisation was applied to the entire 4D dataset by a single multiplicative factor. The statistical analysis of the time-series was carried out using FILM with local autocorrelation correction (Woolrich et al., 2001). Contrasts of interest were the following: i) the stimulus response (easy + hard trials > implicit baseline), ii) the feedback response (correct + error trials > implicit baseline), iii) the hard > easy (stimulus) response, and iv) the error > correct (feedback) responses.

Within each region of interest (ROI) (see section 2.9) and for each contrast of interest, the task-evoked responses were averaged using a weighted sum, separately for each data set. Subsequently, the t-statistics from the 3 T-A and 3 T-B data sets were averaged in order to compare the 7 T data set to a single 3 T data set. For each contrast of interest, differences in t-statistics were evaluated in a 2-way repeated-measures ANOVA with factors ROI group (levels: cortical regions, striatum, & brainstem nuclei) and field strength (levels: 3 T vs. 7 T) using the Python package *pyvttbl*. Planned comparisons for each individual ROI were done by means of a non-parametric two-tailed permutation test (10,000 permutations, custom Python code). No spatial smoothing was applied for the ROI analyses.

At the whole brain level, field-strength differences for each contrast of interest were investigated separately. The resulting t-statistic of the GLM fits for the 3 T-A and 3 T-B data sets were averaged in order to compare the 7 T data set to a single 3 T data set. Task-evoked responses were formally tested by means of a non-parametric one-sample permutation test with 5000 permutations and variance smoothing (5 mm), separately for the positive and negative directions (Randomise, FSL). The family-wise error rate (FWE; p-value threshold = .05) was controlled using the threshold-free cluster enhancement method (TFCE, FSL). Variance smoothing was applied to the t-maps, as per recommendation by FSL for small samples (< 20). The same procedure was applied to test the 3 T-A and 3 T-B data sets separately (Supplementary Fig. 1).

### 2.8. Characterizing temporal SNR

We computed tSNR as the mean of the time series divided by its standard deviation of residual, ongoing fMRI signal fluctuations, after accounting for the evoked responses. The number of volumes (TRs) included in the computation was equalized across data sets for each participant (according to the minimum number of volumes of either the 3 T-A, 3 T-B or 7 T acquisitions within each session). For each participant and data set, the tSNR of each voxel was computed from the pre-processed time series for each session and then averaged across sessions. This resulted in one tSNR volume per participant per data set. Subsequently, the mean tSNR of the 3 T-A and 3 T-B data sets were averaged in order to compare the 7 T data set to a single 3 T data set.

The mean tSNR value within each ROI (see section 2.9) was weighted based on the probability values of the individual voxels. Subsequently, the tSNR values of the 3 T-A and 3 T-B data sets were averaged in order to compare the 7 T data set to a single 3 T data set. Differences in tSNR were evaluated in a 2-way repeated measures ANOVA with factors ROI group (levels: cortical regions, striatum, & brainstem nuclei) and field strength (levels: 3 T vs. 7 T). Planned comparisons for each individual ROI were done by means of a non-parametric two-tailed permutation test (10,000 permutations, custom Python code). No spatial smoothing was applied for the ROI analyses.

At the whole-brain level, the difference in tSNR for the mean 3 T data set was formally compared with the 7 T data set by means of a non-parametric one-sample permutation test with 5000 permutations and variance smoothing (5 mm), separately for the positive and negative directions (Randomise, FSL). The FWE rate (p-value threshold = .05) was controlled using the threshold-free cluster enhancement method (TFCE, FSL). Variance smoothing was applied to the resulting t-maps for the whole-brain analysis of tSNR, as per recommendation by FSL for small samples (< 20).

### 2.9. Defining regions of interest (ROIs)

A substantial body of previous electrophysiology work on variants of our behavioral task has identified neural correlates of different elementary operations entailed in the task in several cortical and subcortical brain regions, specifically: sensory encoding, decision-making, reward anticipation, and reward feedback processing (e.g. (Britten et al, 1992; de Gee et al., 2017; Ding and Gold, 2013; Donner et al., 2007; Lak et al., 2017; Rees et al., 2000; Siegel et al., 2006; Urai et al., 2017). We thus defined a number of ROIs a priori, based on their involvement in the above task-related operations: Dorsal visual cortical regions based on probabilistic maps of visual topography (Wang et al., 2014) included area hMT (hereafter referred to as MT+), V1, V3AB, and intraparietal sulcus regions 0 and 1 (IPS01); the anterior cingulate cortex (ACC) was based on the Harvard-Oxford Cortical Atlas (FSL). The corpus striatum regions included the caudate, putamen and nucleus accumbens (NAc), based on the Oxford-GSK-Imanova Structural–anatomical Striatal Atlas (FSL). Neuromodulatory brainstem nuclei included the VTA (Ballard et al., 2011), SN (Murty et al., 2014), and LC (Keren et al., 2009). Additionally, three ‘ROI groups’ were defined as the average (tSNR or T-statistic) of the following combinations of individual ROIs: V1, V3AB, MT+, IPS01 and ACC (‘cortical regions); caudate, putamen and NAc (‘striatum’); and LC, SN and VTA (‘brainstem nuclei’).

All ROI masks used here were probabilistic. To obtain an unbiased probability value in the combined masks (dorsal and ventral parts of V1, V3A and V3B, and IPS0 and IPS1), the probability values in overlapping voxels were averaged. For all ROIs, we combined homotopic regions in both hemispheres into a single ROI. All ROI masks were restricted spatially by the common field of view between the 3 T and 7 T acquisitions. Finally, within each ROI mask, the probability values were normalized with respect to the maximum probability so that values ranged from 0 to 1. ROI means were weighted based on these normalized probability values (using all voxels within each ROI).

### 2.10 Coherent-motion localizer

We aimed to verify that the atlas-based V3AB and MT+ regions of interest responded significantly to coherent motion as compared with random motion (Braddick et al., 2001; Britten et al., 1993; Newsome and Pare, 1988). To this end, we also ran so-called motion localizer runs, in which blocks of random dot kinematograms at 100% motion coherence were alternated with blocks of random dot kinematograms at 0% coherence. Each block lasted 10 s and showed either coherent dot motion in a single direction or random motion. Eight directions of coherent motion were presented twice each in randomized order (four cardinal and four ordinal directions). In total, 32 blocks were presented per run of the localizer. The order of the coherent and random motion blocks was counterbalanced both within and across participants (i.e. ABAB -> BABA). To control attention and minimize eye movements, participants were instructed to push a button with their right index finger whenever the fixation region changed from grey to green. A change in the fixation region occurred during half of the blocks at random, with the constraint that it occurred an equal number of times during both the coherent motion and random motion blocks. Button-press responses were not analyzed. Within a block, the color-change in fixation occurred at random between one and eight seconds. Motion parameters were the same as indicated for the motion discrimination task at the corresponding MRI field strength with the exception that the dots were presented within the full annulus (not in one half of the visual field). One run of the localizer was administered during each scan session (in total 4 runs for 3 T and 2 runs for 7 T acquisitions) and lasted 5.7 min.

Pre-processing steps for the motion localizer runs were as follows. Each run was (i) brain extracted using the BET, (ii) unwarped based on the session-specific B0 field map, (iii) motion corrected (sinc interpolation, MCFLIRT, FSL), and (iv) high-pass filtered to correct for low frequency drifts (Gaussian-weighted least-squares straight line fitting, window size = 50 samples). For each session, one EPI volume, the motion-corrected mean volume of the third run (out of six) of the 2AFC task, was used as the reference image for all EPI-based registration including motion correction.

The statistical analysis was carried out using FEAT version 6.00 (FSL). At the first-level analysis, two regressors of interest were modeled for each run: coherent and random motion. The contrast of Coherent > Random motion was of interest. Time-series statistical analysis was carried out using FILM with local autocorrelation correction. At the second level, the contrast estimates of two runs from each field-strength acquisition (3 T & 7 T) were averaged for each participant. Statistic images (z-statistic) were thresholded at *p* < .05 uncorrected for multiple comparisons. The mean z-statistic in each ROI were weighted based on the probability values of the ROI masks.

Note that because the visual angles differed between field-strength acquisitions, we could not formally test for differences in the BOLD-acquisition sensitivity of MT+ based on the localizer (Born and Bradley, 2005).

## 3. Results

The current study compared evoked responses of 3 T and 7 T fMRI acquisitions for a number of select brain regions in visual cortex, striatum and brainstem, within the same healthy adult participants. To this end, we measured fMRI responses with near-whole brain coverage at 3 T and 7 T, while participants performed a basic visuo-motor decision task (random dot motion discrimination), followed by the monetary reward-coupled feedback (Fig. 1A). Two difficulty levels were randomly intermixed so that each trial was either hard (70% correct) or easy (85%), depending on individual threshold motion coherence levels that were established in a behavioral session before scanning began (Fig. 1B). The task entailed several operations with well-characterized neural substrates (Gold and Shadlen, 2007): encoding of visual motion information, perceptual decision-making, decision confidence and/or reward anticipation during and right after decision formation; and processing of reward prediction errors upon the delivery of decision outcome in terms of feedback tones.

**Fig. 1.**
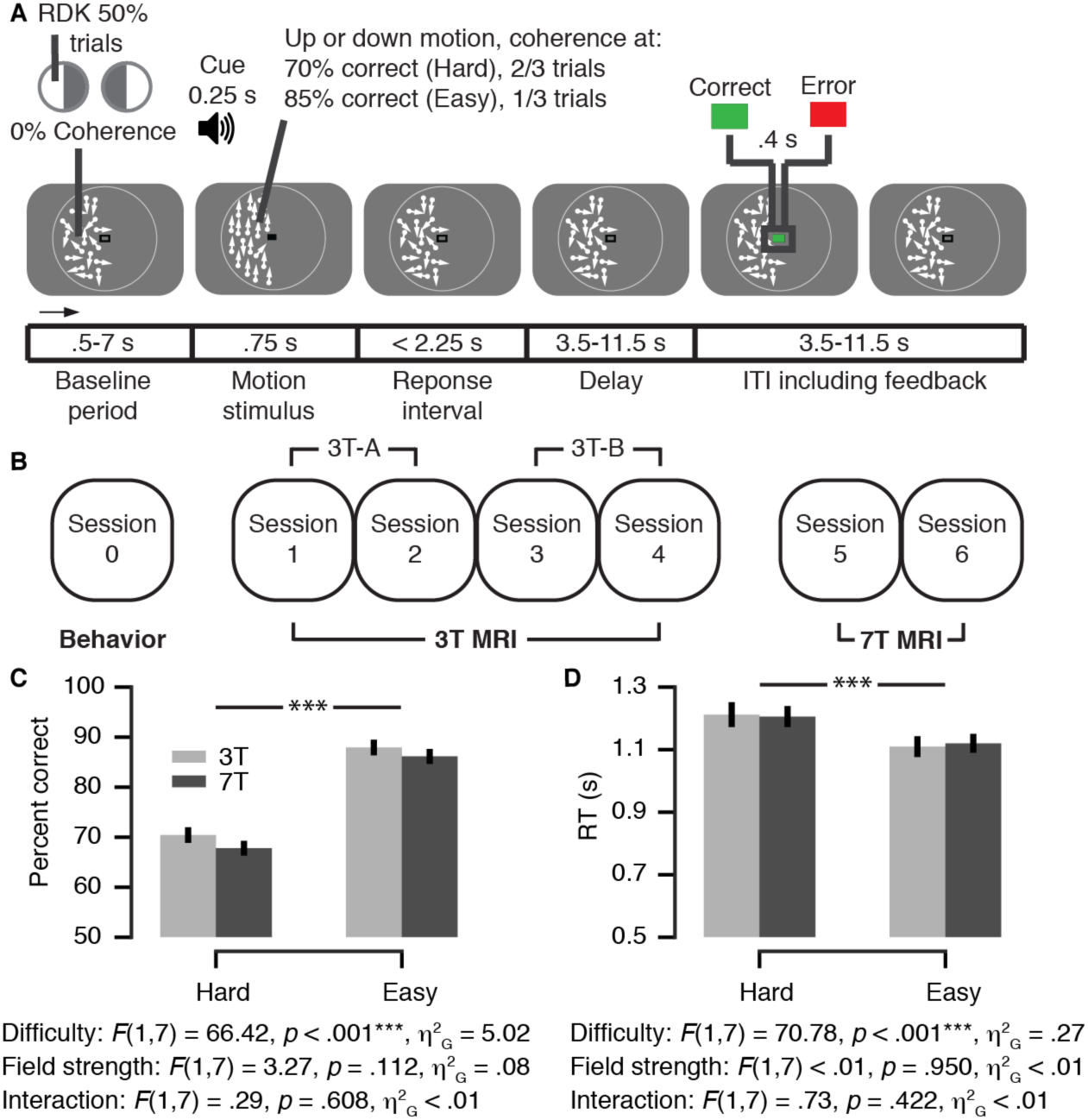
Experimental design and behavior. (**A**) Behavioral task. Subjects performed a 2AFC motion direction discrimination task during fMRI. Random dot kinematograms (RDK) were shown in one visual hemifield on each run (hemifields alternating sides across runs). The decision interval ranged from onset of the motion stimulus to the participant’s response. The feedback interval ranged from feedback onset into the subsequent inter-trial interval (ITI). Feedback was coupled to a monetary reward based on the accuracy of each run of 25 trials. (**B**) Experimental protocol. Participants completed seven sessions in total, including four 3 T and two 7 T MRI sessions. The data of the two 7 T sessions were averaged. The 3 T MRI data was split into two subsets, 3 T-A and 3 T-B, and averaged (i.e. ‘3 T’) in order to compare the 7 T to a single data set. (**C, D**) Choice accuracy (C) and mean RT (D) as a function of task difficulty and field strength (repeated measures 2-way ANOVA). F-statistics, main effect of task difficulty, field strength, and their interaction. Error bars, s.e.m. (*N* = 8). Panel (A) is reproduced from (Colizoli et al., 2018).

All participants in the present analysis took part in four 3 T MRI sessions and then two 7 T MRI sessions (Fig. 1B). Subjects’ choice behavior was faster and more accurate in the easy compared to the hard condition of the discrimination task (Fig. 1C,D). Critically, neither RTs nor accuracy differed between the two scanner environments (Fig. 1C,D main effect of field strength) and there was no significant interaction between task difficulty and field strength (Fig. 1C,D).

### 3.1 Comparison of evoked responses at 7 T and 3 T for different brain regions

We analyzed evoked fMRI responses of a number of cortical and subcortical brain regions, which are implicated in different operations engaged in the visual motion discrimination task (sensory encoding, decision-making, decision certainty / reward anticipation, feedback processing) (Britten et al., 1992; de Gee et al., 2017; Ding and Gold, 2013; Donner et al., 2007; Lak et al., 2017; Rees et al., 2000; Siegel et al., 2006; Urai et al., 2017): (i) cortex: three (clusters of) cortical areas along the dorsal pathway analyzing visual motion (V1, V3AB, MT+) or involved in task/decision processing (IPS01 and ACC); (ii) the corpus striatum (caudate nucleus, nucleus accumbens, putamen); and (iii) brainstem: three modulatory nuclei in the pons midbrain, all of which contribute to decision-making, but whose activity is notoriously difficult to measure at 3 T: the noradrenergic LC and the dopaminergic SN and VTA.

We first verified that an established pattern of sensory responses in visual cortex was present in our data. In separate localizer runs (see Methods), we found that extrastriate visual cortical regions V3AB and MT+, but not V1, exhibited the sensory responses specific to coherent compared to incoherent dot motion expected from previous work (Huk et al., 2002; Tootell et al., 1995, 1997). This was true and robust for both field strengths (Fig. 2; all p-values < .05). Please note that the voxel size and the extent of the visual stimulus were both smaller at 7 T than 3 T (both factor of ∼1.8, see Methods). Smaller voxels and smaller stimuli (driving a smaller portion of the retinopically organized visual cortical areas (Huk and Heeger, 2002; Wang et al., 2014) are both expected to reduce overall fMRI responses in these cortical regions. Nonetheless, we found about equally strong (statistically indistinguishable, all p-values > .2) responses in both V3AB and MT+. During the motion discrimination task, we also found high across-session overlap of the responses in visual cortex at 3 T, which were performed in four different sessions (Supplementary Fig. 1).

**Fig. 2.**
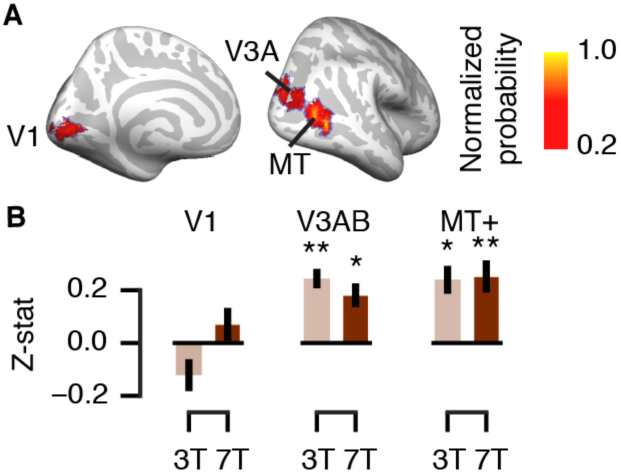
Visual cortical responses to coherent vs. incoherent random dot motion (localizer runs). (**A**) Regions of interest. Probabilistic mask from Wang et al. (2014) for visual cortical areas V1, V3AB and MT+ displayed on the cortical surface, thresholded at *p* = .2 for illustration. Each mask was normalized with respect to its maximum value, yielding a normalized probability. (**B**) Differential responses of each area to 100% coherent relative to incoherent (0%) dot motion. \**p* < .05, \*\**p* < .01 (two-tailed permutation tests against zero). Error bars, s.e.m. (*N* = 8).

Having thus established the quality of our data, we grouped the brain regions into three groups listed above (cortical regions, striatum, and brainstem nuclei), for which we expected differential BOLD-acquisition sensitivity on fMRI field strength. Please note that each of these groups consisted of precisely delineated brain regions, some of which were small (the brainstem nuclei). For each of these groups we then quantified the task-evoked responses as t-statistics (i) during the decision interval of the task (i.e., time-locked locked to the onset of the dot-motion stimulus; see Methods); and (ii) the post-feedback interval, time-locked to the feedback color (green or red) that informed subjects about the correctness of their preceding choice (Fig. 1A). For each interval, we quantified two distinct components of the response: the overall response (compared to pre-trial baseline, left panels for each interval) or the response difference between hard and easy decisions (decision interval) and between error and correct feedback.

For the overall response during the decision interval, we observed a significant interaction between ROI group and fMRI field strength (Fig. 3B, see Supplementary Fig. 2 for the individual cortical and striatum ROIs). Hereby, the biggest improvement at 7 T seemed to occur for the brainstem nuclei and smallest improvement for the cortical regions. Similar trends were evident, but not statistically significant, for the differential responses (i.e. hard vs. easy and error vs. correct) in the decision and feedback intervals (Fig. 3B).

**Fig. 3.**
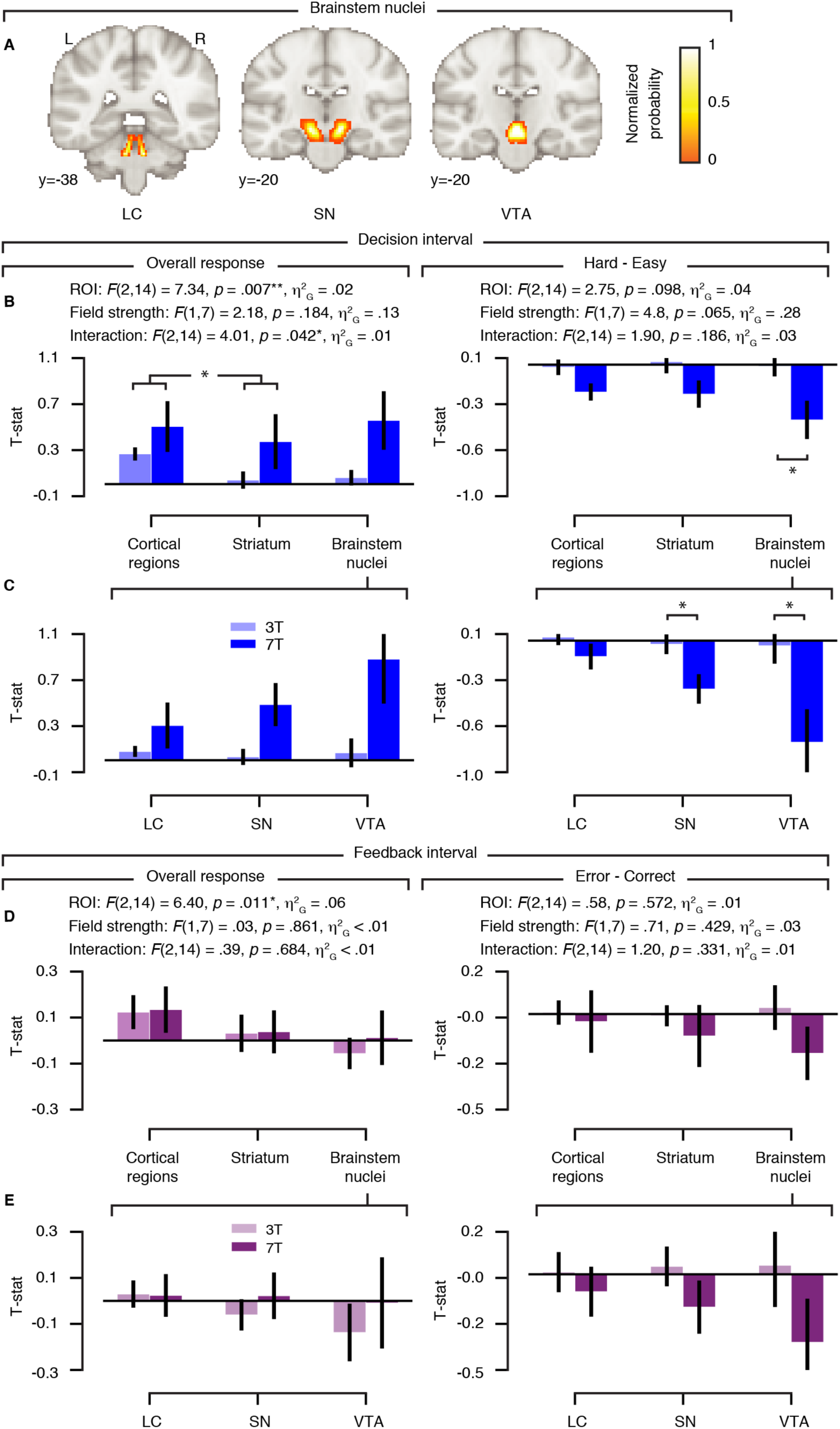
Task-evoked responses for 3 T and 7 T acquisition in different brain regions. (**A**) Brainstem and masks are illustrated on the MNI152 brain. Each probabilistic mask (see Methods) was normalized with respect to its maximum value. LC, locus coeruleus. SN, substantia nigra. VTA, ventral tegmental area. (**B**) Amplitude of evoked response per ROI group (cortical regions, striatum & brainstem nuclei) and field strength for the decision interval (stimulus locked). (**C**) Planned comparisons within the brainstem nuclei (decision interval). (**D**) Amplitude of evoked response per ROI group and field strength for the post-feedback interval. (**E**) Planned comparisons within the brainstem nuclei (post-feedback interval). (B-E) The left column shows the overall response (compared to pre-trial baseline); the right column shows the amplitude difference between hard vs. easy trials (decision interval, blue) and error vs. correct feedback (post-feedback interval). 3 T amplitudes were averaged across 3 T-A and 3 T-B data subsets. F-statistics, main effect of ROI group, field strength, and their interaction. \**p* < .05, \*\**p* < .01 (two-tailed permutation tests). Error bars, s.e.m. (*N* = 8).

A significant main effect of ROI group was observed for the overall responses during both decision and feedback intervals. The main effect of field strength was not statistically significant, but showed a trend for the differential responses (hard vs. easy) during the decision interval (*p* = .065).

Similar trends were evident for all individual brainstem ROIs (Fig. 3C), which suggests a particular benefit from higher field strength. In particular, evoked differential responses during the decision interval were significantly stronger at 7 T than at 3 T in the VTA and SN (all p-values < .05).

No significant differences between field strengths for the task-evoked responses were obtained at the level of the whole brain.

### 3.2 Comparison of temporal SNR at 7 T and 3 T for different brain regions

Our current study focused on the comparison of field-strength dependence of evoked fMRI responses between different brain regions: in particular brainstem nuclei versus larger, easier to image structures, such as striatum and cortex. This focus was motivated by strong prior hypotheses regarding these responses across brain regions. Another measure, temporal signal-to-noise-ratio (tSNR), is commonly used as a proxy for fMRI image quality (Triantafyllou et al., 2005; Vu et al., 2017) and has frequently been compared between field strengths (Brooks et al., 2013). By dividing the temporal mean of the fMRI signal by its temporal variability across a run, this measure quantifies the stability of the signal (albeit being affected by different noise sources, see Discussion). We also compared tSNR between the 3 T and 7 T acquisitions in an additional analysis. To this end, we used the residual signal fluctuations remaining after fitting the GLM used for quantifying the evoked responses described in the previous section (see Methods for details). We analyzed tSNR across the brain, at the single-voxel level, and at the level of the same ROIs for which we evaluated the evoked responses.

As for the analysis of evoked responses, we averaged tSNR values of the 3 T-A and 3 T-B data sets (i.e. ‘3 T’) before comparison with the 7 T data set. No significant difference in tSNR between field strengths was present in any region at the single-voxel level (Fig. 4). Again, please note that the 7 T voxels were 1.8-fold smaller as compared with the 3 T acquisition, still leading to about the same tSNR.

**Fig. 4.**
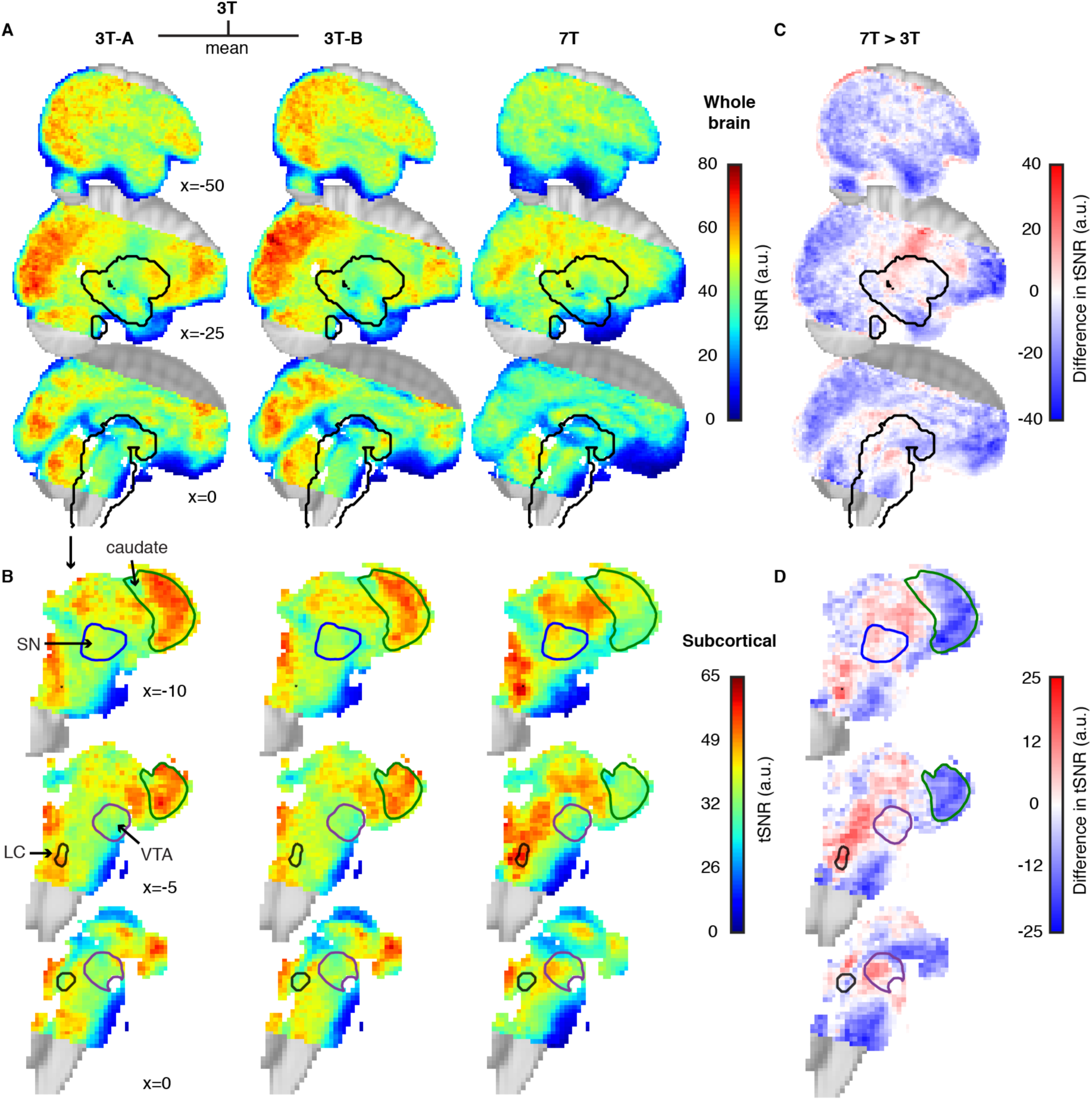
Spatial distribution of the temporal signal-to-noise ratio (tSNR) during the 2AFC task. (**A**) tSNR distributions for the 3 T-A, 3 T-B and 7 T data sets at the whole-brain level (*N* = 8) and (**B**) subcortical partition of the brain volume (black outlines in panel A) partition extracted and magnified. The tSNR of each data set was computed with an equal number of units (TRs) per participant. (**C**) Group average tSNR of the 7 T data set compared to the mean of the two 3 T data sets (3 T-A and 3 T-B) at the whole-brain level and (**D**) subcortical partition extracted and magnified. Structures are labeled based on probabilistic atlases, including the caudate, locus coeruleus (LC), substantia nigra (SN) and ventral tegmental area (VTA). Colored regions indicate the common field of view of the 3 T and 7 T acquisitions (MNI152 brain as background).

At the level of ROIs, we again found a significant interaction between ROI group and field strength (Fig. 5A). A main effect of ROI group was observed. The main effect of field strength was not significantly significant. Just as for the evoked responses in Fig. 3 (decision interval), tSNR was higher in the brainstem ROI group, but not for the cortical regions and striatum, for the 7 T as compared with the 3 T acquisition (see Supplementary Fig. 3 for the individual ROIs).

**Fig. 5.**
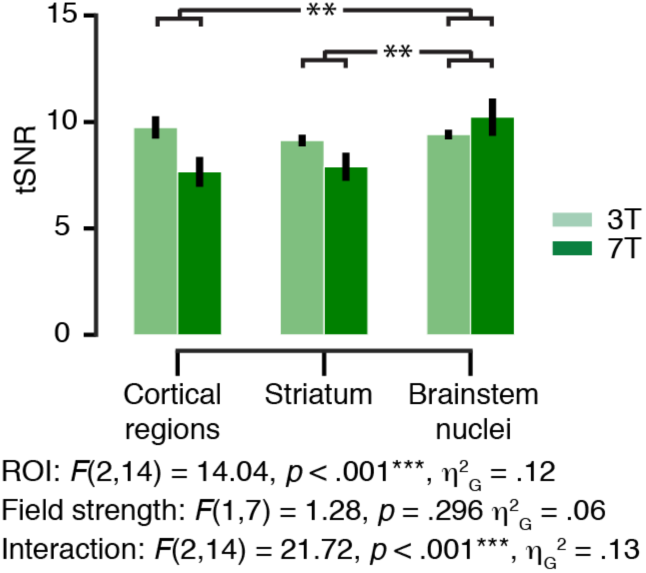
Region of interest (ROI) analysis of the temporal signal-to-noise ratio (tSNR) during the motion discrimination task. TSNR for the 3 T and 7 T acquisitions for each ROI group (cortical regions, striatum & brainstem nuclei). The mean tSNR of each data set was computed with an equal number of units (TRs) per participant. The 3 T data here are the mean of the 3 T-A and 3 T-B data subsets. F-statistics, main effect of ROI group, field strength, and their interaction. \*\**p* < .01 (two-tailed permutation tests). Error bars, s.e.m. (*N* = 8).

No significant differences between field strengths in tSNR were obtained at the level of the whole brain.

## Discussion

We used an extensively studied cognitive task (visual perceptual decisions under uncertainty) to perform a systematic, within-subject, head-to-head comparison of evoked responses between 3 T and 7 T fMRI acquisitions, across a range of well-defined cortical and subcortical brain regions. Overall, our findings are consistent with the notion that sensitivity for evoked responses in the small brainstem nuclei (during the decision interval and to a lesser extent also post-feedback interval) benefitted most from 7 T measurements.

We did not opt for an exact match of imaging parameters, in particular spatial resolution, between field strengths, but rather chose resolutions that would be common choices for each field strength (e.g. (Hale et al., 2010; Morris et al., 2019; Theysohn et al., 2013; Torrisi et al., 2018). Accordingly, spatial resolution was higher for our 7 T than 3 T measurements (see Methods). Nonetheless, we obtained equally strong, or even stronger responses at 7 T, in particular in brainstem regions. This highlights the utility of 7 T measurements for high-precision fMRI, in particular in brainstem structures, the activity of which has been notoriously difficult to monitor with 3 T fMRI.

Other imaging parameters apart from spatial resolution also differed between our 3 T and 7 T measurements, but we consider it unlikely that these differences have biased our current findings. The used echo time (TE) at 3 T was somewhat longer than 7 T: still both TEs were rather short compared to the cortical gray matter T2* (Peters et al., 2007), especially at 3 T. This allows for whole-brain coverage and a reasonable TR and is wide-spread practice in cognitive neuroimaging. The SENSE factor, bandwidth and EPI factor contribute to differences in the amount of susceptibility induced image distortion, but these differences are much smaller than those caused by the B0 difference. Echo planar images at 7 T are more distorted than those at 3 T, but both are corrected by B0-field unwarping and therefore should not have affected the BOLD measures. Finally, also the radio frequency (RF) coils differed between the 3 T and 7 T measurements. Indeed, the properties of the RF coil significantly influence the SNR distribution over the brain (Wiggins et al., 2006). Yet, we used two state-of-the-art, vendor-provided coils offering whole-head coverage.

The main interest of cognitive neuroimaging in 7 T fMRI lies in the ability to detect small-amplitude variations of brain activity even at high spatial resolution (Welvaert and Rosseel, 2013). For example, (van der Zwaag et al., 2009) found that the number of activated voxels, t-statistics, and percent signal change all increased significantly with field strength (1.5T, 3 T and 7 T) in the same common ROI in a head-to-head comparison of six participants performing a simple motor task. Stronger bilateral hippocampus activation during memory encoding was obtained at 7 T fMRI as compared with 3 T (Theysohn et al., 2013). More recently, (Torrisi et al., 2018) concluded that 7 T fMRI resulted in significant gains compared to 3 T in terms of effect size, detection of smaller effects, and group-level power while measuring response inhibition (a standard GO / NOGO task). To our knowledge, the current study is the first that directly compared the benefits of 7 T fMRI field strength between cortical, striatal, and brainstem regions.

In line with our findings, (de Hollander et al., 2017) stress that common acquisition parameters (at higher field-strength) may be optimized for specific brain regions and furthermore that brain regions differ in terms of physiological noise contributions and distance from the receiver coils. The authors did not find reliable hemodynamic responses in the brainstem’s iron-rich basal ganglia nuclei during a robust stop-signal reaction task using ‘standard’ 7T imaging parameters. Successfully imaging the basal ganglia nuclei at 7T required a shorter TE (∼15 ms) as compared to cortex (25-35 ms), but also other factors interacted with signal detectability, for instance, the spatial resolution and acceleration factor.

Although we found both higher tSNR and stronger task-evoked responses in the brainstem nuclei, we are cautious with our interpretation of a (causal) relationship. The relationship between tSNR and BOLD-acquisition sensitivity is complex and non-linear (Murphy et al., 2007; Triantafyllou et al., 2005; Wald, 2012; Welvaert and Rosseel, 2013). Temporal SNR combines non-neural (physiological and thermal) noise and genuine, ongoing neural variability (Fox and Raichle, 2007), which is also abundant in electrophysiological measurements of neural mass action (Hipp et al., 2012). Hence, we here focused on comparing the amplitude of evoked fMRI responses between field strengths, in a well-controlled task design with established neurophysiological underpinnings.

The differential responses in brainstem (in particular VTA/SN) during decision intervals we observed here were largely in line with a coding of confidence and/or reward anticipation, as observed in previous single-unit recordings from dopaminergic midbrain neurons in monkeys (Lak et al., 2017). Likewise, the differential responses to the feedback are in line with coding of reward prediction errors, also in line with (Lak et al., 2017). Thus, our findings add to a growing body of literature implicating dopamine in perceptual decision-making under uncertainty (de Lafuente and Romo, 2011; Lak et al., 2017; Nomoto et al., 2010). Specifically, dopaminergic midbrain neurons encode not only prediction errors upon feedback about decision outcome, but they encode a graded ‘belief state’ about the accuracy of a choice, given the evidence and choice (Lak et al., 2017); for indirect measurements in humans see (Colizoli et al., 2018; Urai et al., 2017). This graded belief state is analogous with a graded reward anticipation that evolves after (or even during) decision formation and is maintained until the delivery of feedback, evident in brainstem activity (Lak et al., 2017) as well as behavioral proxy variables (Colizoli et al., 2018; Urai et al., 2017).

In conclusion, our results highlight the potential of 7 T fMRI for illuminating the contribution of subcortical structures, in particular small neuromodulatory brainstem nuclei such as the VTA, to cognition in the human brain. Computational theory (Dayan, 2012; Montague et al., 2004) and physiological evidence (e.g., (Aston-Jones and Cohen, 2005; D’Ardenne et al., 2008; de Gee et al., 2017; Engelhard et al., 2019; Iglesias et al., 2013; Lak et al., 2017; Varazzani et al., 2015) point to a key role of these brainstem structures in the orchestration of cognitive computations operating in higher-tier brain regions. Yet, progress in pinpointing their computational role, and in particular, their dynamic interaction with the cortical networks implementing cognitive computations, has so far been largely limited by the technical challenges entailed in monitoring the activity of small brainstem nuclei such as the VTA with the necessary spatial precision and sensitivity. We envisage that the progressive establishment of 7 T fMRI in neuroimaging centers will lead to a surge of studies illuminating the role of the brainstem in cognition. This will not only deepen insight into healthy cognition, but also advance the mechanistic understanding of psychiatric disorders (Montague et al., 2004; Stephan et al., 2015)

## Abbreviations

T: Tesla
MRI: magnetic resonance imaging
fMRI: functional magnetic resonance imaging
SNR: signal-to-noise ratio
tSNR: temporal signal-to-noise ratio
BOLD: blood oxygenation level dependent
HRF: hemodynamic response function
RETROICOR: retrospective image correction technique
ROI: region of interest
2AFC: two-alternative forced choice
EPI: echo-planar images
RF: radio frequency
TFCE: threshold-free cluster enhancement
FWE: family-wise error rate
FSL: fMRIB software library
MNI: Montreal Neurological Institute
V1: primary visual cortex
V3AB: third visual cortex regions A and B
MT+: human medial temporal area
IPS01: intraparietal sulcus regions 0 and 1
ACC: anterior cingulate cortex
NAc: nucleus accumbens
LC: locus coeruleus
SN: substantia nigra
VTA: ventral tegmental area.

## Acknowledgments

This research was funded by the European Union Seventh Framework Programme (FP7/2007-2013) under grant agreement no. 604102 (Human Brain Project) (to T.H.D.), and the Deutsche Forschungsgemeinschaft (DFG, German Research Foundation) DO 1240/3-1, DO 1240/2-1, and SFB 936/A7 (to T.H.D.). This work was carried out on the Dutch national e-infrastructure with the support of the SURF Cooperative (www.surfsara.nl). We thank Tomas Knapen for contributing analysis materials, Birte U. Forstmann and Max C. Keuken for advice on the experimental design.

## Author contributions

O.C. contributed Conceptualization, Investigation, Formal Analysis, Analysis Materials, Writing - original draft, Writing – review and editing; J.W.d.G. contributed Conceptualization, Investigation, Formal Analysis, Analysis Materials, Writing - review and editing; W.v.d.Z contributed Methodology, Investigation, Writing - review and editing; T.H.D. contributed Conceptualization, Resources, Supervision, Writing - original draft, Writing - review and editing.

## Competing interests

The authors declare no competing interests.

## Data and code availability

The data and analysis code will be made publicly available upon publication (DOI:xxx) and comply with the funding bodies requirements.

## Materials & Correspondence

(O.C.) and (T.H.D.)

## Supplementary materials

**Supplementary Fig. 1.**
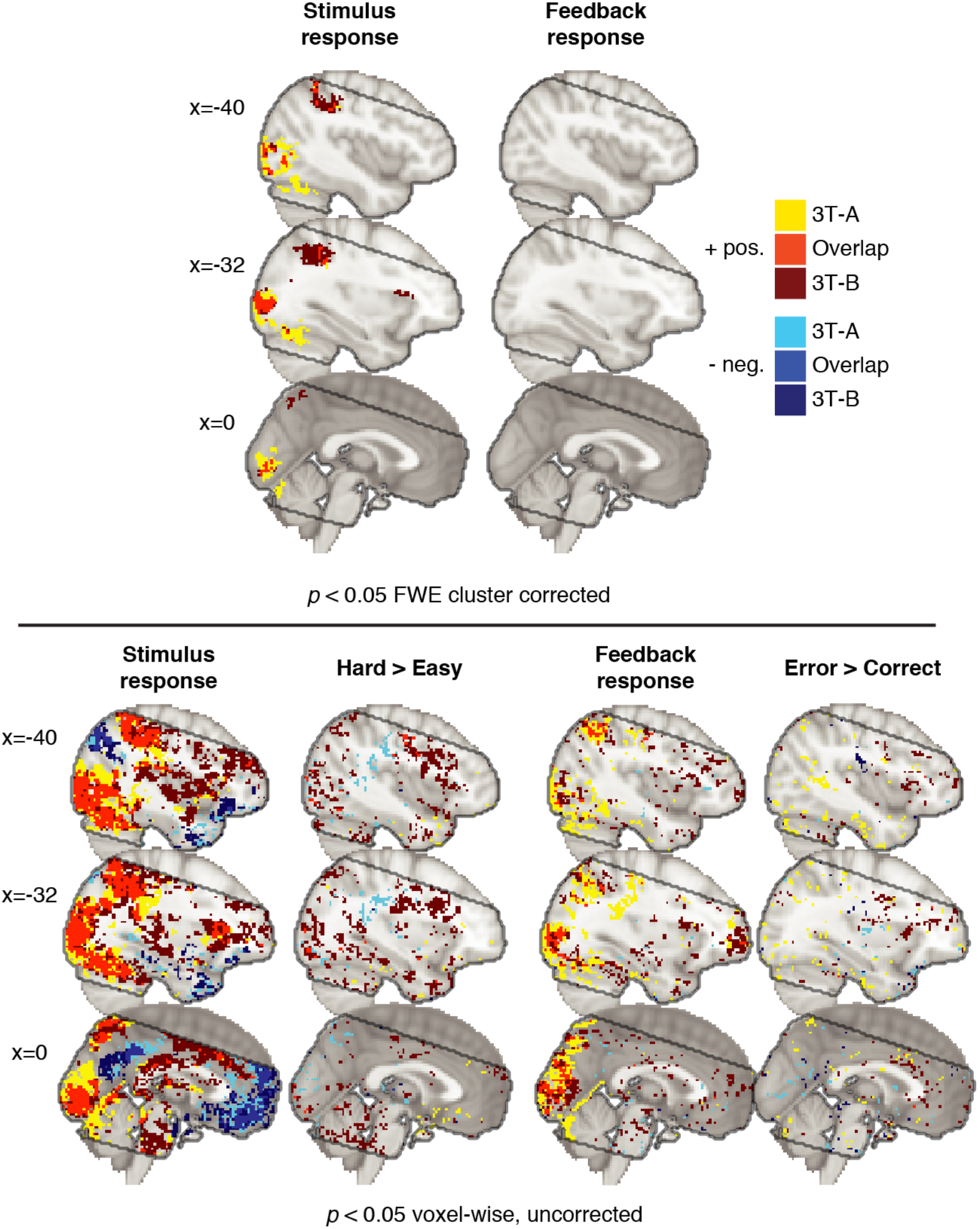
Whole brain results of the task-evoked responses for the 3 T acquisition. Contrasts of interest were investigated in a GLM analysis with an equal number of trials for each data set per participant (*N* = 8). (Top) Voxels surviving cluster-correction are shown for the 3 T-A and 3 T-B data sets for the stimulus and feedback responses, from the decision and post-feedback interval, respectively (*p* < .05, FWE, corrected). (Bottom) Voxels significant before cluster-correction for all contrasts of interest (*p* < .05, voxel-wise, uncorrected). The grey outline indicates the common field of view of the 3 T and 7 T acquisitions (MNI152 brain as background).

**Supplementary Fig. 2.**
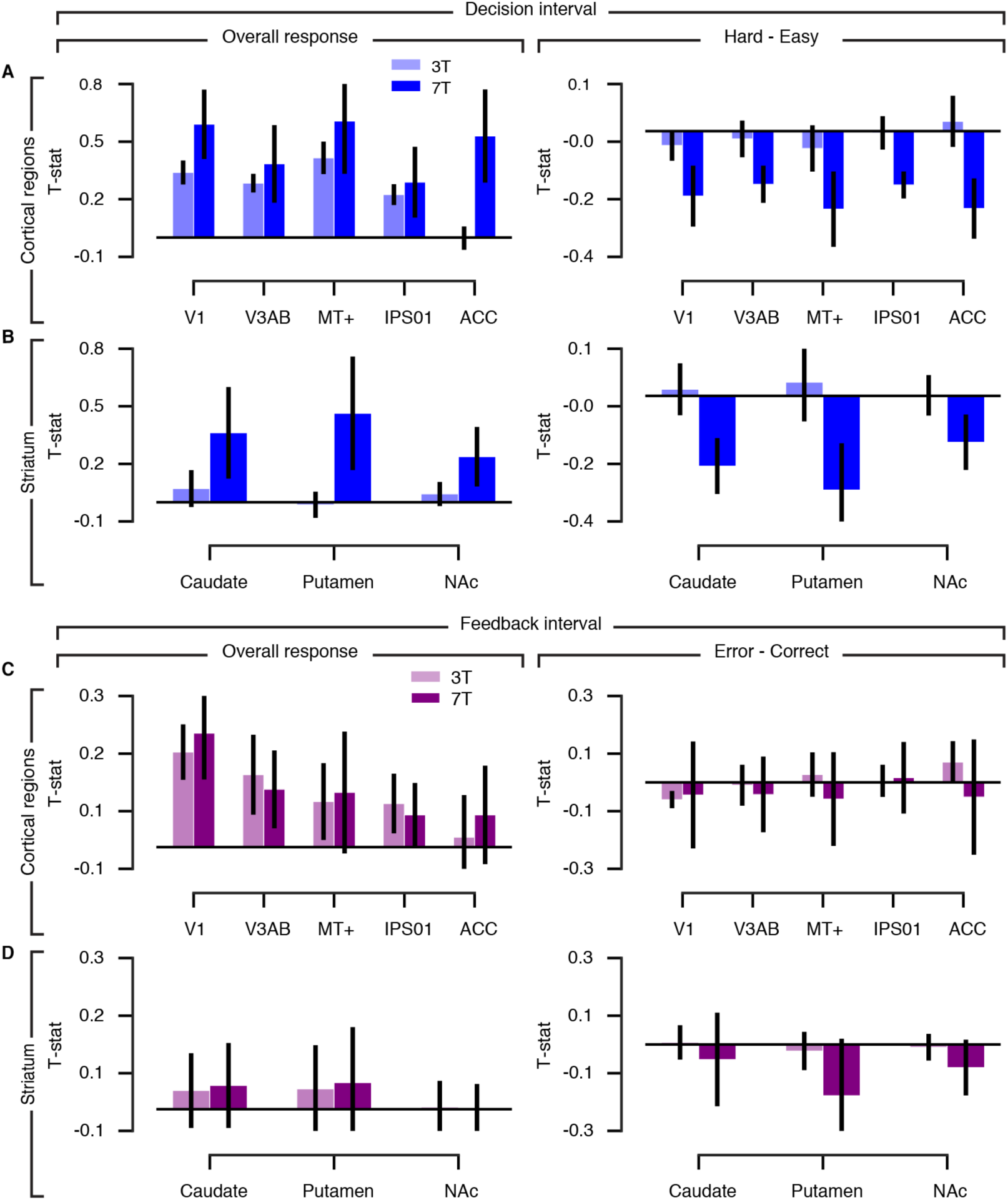
Task-evoked responses for 3 T and 7 T acquisition in cortical regions and striatum. (**A, B**) For the decision interval (stimulus-locked), the overall response and amplitude differences between hard vs. easy trials were investigated in cortical regions (A) and the striatum (B). NAc, nucleus accumbens. (**C, D**) For the post-feedback interval, the overall response and amplitude differences between error vs. correct trials were investigated in cortical regions (C) and the striatum (D). No field-strength differences were obtained (two-tailed permutation tests). The 3 T data are the mean of the 3 T-A and 3 T-B data subsets. Error bars, s.e.m. (*N* = 8).

**Supplementary Fig. 3.**
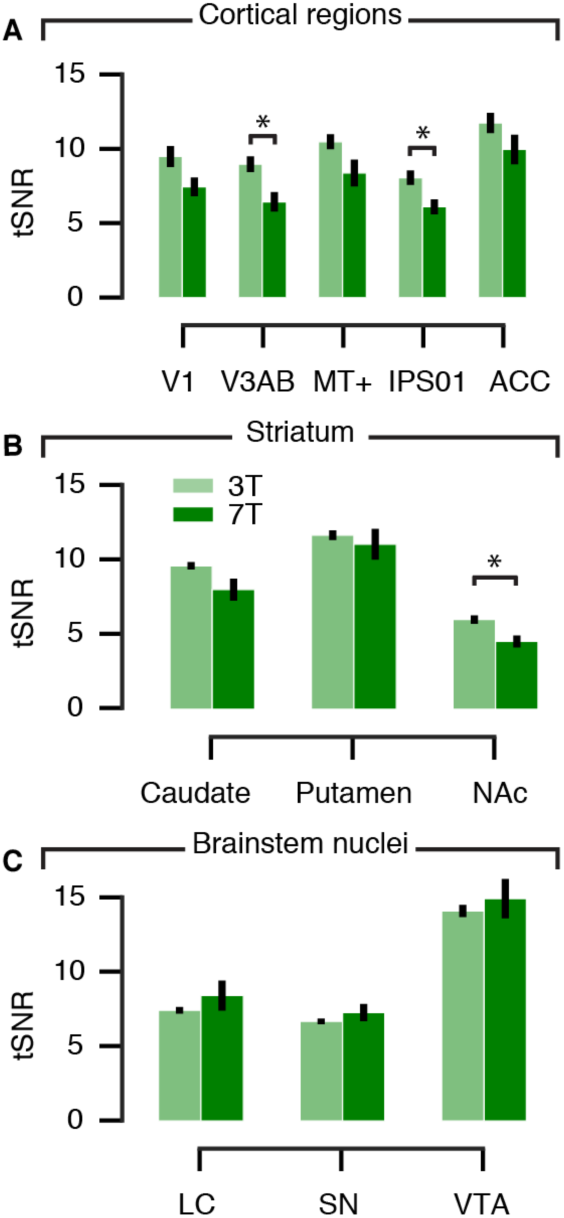
Temporal signal-to-noise ratio (tSNR) for 3 T and 7 T acquisition in cortical regions, striatum and brainstem nuclei. (**A**) Cortical regions (**B**) Striatum. NAc, nucleus accumbens. (**C**) Brainstem nuclei. LC, locus coeruleus. SN, substantia nigra. VTA, ventral tegmental area. The mean tSNR of each data set was computed with an equal number of units (TRs) per participant. The 3 T data here are the mean of the 3 T-A and 3 T-B data subsets. \**p* < .05 (two-tailed permutation tests). Error bars, s.e.m. (*N* = 8).

## Notes

### Competing Interest Statement

The authors have declared no competing interest.

